## Supplemental Tables 1-6 for "CNS myelination requires VAMP2/3-mediated membrane expansion in oligodendrocytes"

| vSNARE | **VAMP2-pHluorin** | | | | | | **VAMP3-pHluorin** | | | | | |
|---|---|---|---|---|---|---|---|---|---|---|---|---|
| Differentiation stage | OPC | | pre-myelinating | | mature | | OPC | | pre-myelinating | | mature | |
| Subcellular location | soma | process | soma | process | soma | sheet | soma | process | soma | process | soma | sheet |
| Mean (events/cell/min) | 1.62 | 3.62 | 1.08 | 6.53 | 0.712 | 7.82 | 1.58 | 5.21 | 0.886 | 9.49 | 0.844 | 11.6 |
| Std. Error of Mean | 0.347 | 0.737 | 0.424 | 2.1 | 0.118 | 1.34 | 0.248 | 0.644 | 0.414 | 0.721 | 0.228 | 2.1 |
| Lower 95% CI | 0.129 | 0.452 | -0.741 | -2.5 | 0.203 | 2.05 | 0.516 | 2.44 | -0.896 | 6.39 | -0.137 | 2.6 |
| Upper 95% CI | 3.12 | 6.79 | 2.9 | 15.6 | 1.22 | 13.6 | 2.65 | 7.98 | 2.67 | 12.6 | 1.82 | 20.7 |
| # of cells (n) | 9 | | 27 | | 22 | | 13 | | 22 | | 34 | |

**Supplementary Table 1:** Descriptive statistics for exocytotic events in cultured oligodendrocytes (CI = confidence interval)

| vSNARE | **VAMP2-pHluorin** | | | | | | **VAMP3-pHluorin** | | | | | |
|---|---|---|---|---|---|---|---|---|---|---|---|---|
| Differentiation | OPC/pre-myelinating | | early myelinating | | mature | | OPC/pre-myelinating | | early myelinating | | mature | |
| Subcellular location | soma | process | soma | sheath | soma | sheath | soma | process | soma | sheath | soma | sheath |
| Mean (events/cell/min) | 0.0396 | 0.621 | 0.0461 | 0.393 | 0.0592 | 0.162 | 0.0943 | 1.27 | 0.193 | 0.649 | 0.159 | 0.364 |
| Std. Error of Mean | 0.0254 | 0.249 | 0.0358 | 0.137 | 0.0289 | 0.0426 | 0.0363 | 0.135 | 0.081 | 0.306 | 0.101 | 0.0567 |
| Lower 95% CI | -0.0412 | -0.172 | -0.0533 | 0.0126 | -0.00917 | 0.0614 | 0.0107 | 0.955 | -0.836 | -3.24 | -0.0886 | 0.226 |
| Upper 95% CI | 0.12 | 1.41 | 0.145 | 0.773 | 0.128 | 0.263 | 0.178 | 1.58 | 1.22 | 4.53 | 0.407 | 0.503 |
| # of cells (n) | 4 | | 5 | | 8 | | 9 | | 2 | | 7 | |

**Supplementary Table 2:** Descriptive statistics for exocytotic events in zebrafish spinal cords (CI = confidence interval)

| | **Vamp2** | | **Vamp3** | |
|---|---|---|---|---|
| **Total events within myelin sheaths** | 31 | | 45 | |
| **# of cells** | 10 cells (in 8 animals) | | 8 (in 7 animals) | |
| **# of sheaths** | 25 (out of 261) | | 25 (out of 158) | |
| **Total length of sheaths (μm)** | 421.2 | | 597.8 | |
| **Total length of paranodes (μm)** | 143.6 | | 146.5 | |
| **Total length of internodes (μm)** | 277.6 | | 451.3 | |
| | **paranode** | **internode** | **paranode** | **internode** |
| **%length (predicted frequency)** | 34.1 | 65.9 | 24.5 | 75.5 |
| **%events (measured frequency, mean ± SEM)** | 50 ± 22.4 | 50 ± 22.4 | 24.9 ± 12.7 | 75.1 ± 12.7 |

**Supplementary Table 3**: Spatial frequency of exocytosis in myelin sheaths

| Day in differentiation media | 3 | | 5 | | 7 | |
|---|---|---|---|---|---|---|
| genotype | **ctrl** | **iBot** | **ctrl** | **iBot** | **ctrl** | **iBot** |
| Mean surface area ($\mu m^2$) | 6099 | 5047 | 12022 | 3557 | 16261 | 8735 |
| Std. Error of Mean | 595 | 240 | 1731 | 325 | 849 | 1144 |
| Lower 95% CI | 4448 | 4380 | 7572 | 2656 | 13904 | 5560 |
| Upper 95% CI | 7750 | 5714 | 16471 | 4458 | 18618 | 11911 |
| total # of cells from n = 5 biological replicates | 310 | 256 | 344 | 254 | 282 | 252 |

**Supplementary Table 4:** Descriptive statistics for membrane surface area of primary oligodendrocytes in culture (CI = confidence interval)

| Gene | Description | control biotinylated - rep1 | control biotinylated - rep2 | iBot biotinylated - rep1 | iBot biotinylated - rep2 | iBot biotinylated - rep3 | non-biotinylated - rep1 | non-biotinylated - rep2 | non-biotinylated - rep3 | log2(fold enrichment) | q-value (adjusted p-value) |
|---|---|---|---|---|---|---|---|---|---|---|---|
| Immt | MIC complex subunit Mic60 | 59 | 19 | 19 | 7 | NaN | NaN | NaN | NaN | -6.25 | 0.050 |
| Nfasc | Neurofascin | 27 | 10 | NA | NaN | NaN | NaN | NaN | NaN | -4.90 | 0.049 |
| Cntn1 | Contactin-1 | 51 | 18 | 7 | 3 | NaN | NaN | NaN | NaN | -4.43 | 0.052 |
| Atp5f1b | ATP synthase subunit beta, mitochondrial | 124 | 36 | 56 | 16 | 24 | NaN | NaN | 1 | -4.40 | 0.048 |
| Mbp | Isoform 4 of Myelin basic protein | 77 | 48 | 21 | 4 | 3 | 6 | 7 | 15 | -4.05 | 0.048 |
| Ank3 (AnkG) | Ankyrin-3 (also Ankyrin G) | 12 | 9 | NA | 2 | NaN | NaN | 2 | 3 | -3.82 | 0.053 |
| Hspa2 | Heat shock-related 70 kDa protein 2 | 9 | 9 | NA | 3 | 1 | NaN | 3 | 5 | -3.25 | 0.046 |
| Rtn4 | Reticulon-4 | 88 | 62 | 9 | 6 | 3 | 5 | 17 | 12 | -3.18 | 0.052 |
| Tppp | Tubulin polymerization-promoting protein | 10 | 8 | 4 | 1 | 1 | NaN | 3 | NaN | -3.01 | 0.046 |
| Mag | Myelin-associated glycoprotein | 13 | 20 | 6 | 2 | 1 | NaN | 2 | 2 | -2.98 | 0.049 |
| Ndufa10 | NADH dehydrogenase 1 alpha subcomplex subunit 10, mitochondrial | 5 | 6 | NA | NaN | NaN | NaN | NaN | NaN | -2.97 | 0.055 |
| Bin1 | Myc box-dependent-interacting protein 1 | 7 | 10 | 2 | NaN | 1 | NaN | NaN | NaN | -2.90 | 0.048 |
| Sept8 | Septin-8 | 10 | 6 | NA | 2 | 2 | NaN | NaN | 4 | -2.81 | 0.045 |
| Dst | Dystonin | 52 | 52 | NA | NaN | NaN | NaN | NaN | 16 | -4.70 | 0.054 |
| Tns3 | Tensin-3 | 14 | 28 | NA | 1 | NaN | NaN | NaN | 7 | -3.91 | 0.048 |
| Frmd4a | FERM domain-containing protein 4A | 13 | 7 | NA | NaN | NaN | NaN | 3 | NaN | -3.82 | 0.054 |
| Capzb | F-actin-capping protein subunit beta | 6 | 4 | NA | NaN | NaN | NaN | NaN | NaN | -2.78 | 0.045 |
| Rhoa | Transforming protein RhoA | 11 | 20 | NA | 2 | 2 | 3 | 3 | 7 | -2.60 | 0.050 |
| Dctn4 | Dynactin subunit 4 | 6 | 5 | NA | 2 | NaN | NaN | 2 | 2 | -2.50 | 0.047 |
| Cdk5 | Cyclin-dependent-like kinase 5 | 3 | 3 | NA | 1 | NaN | NaN | 2 | NaN | -2.49 | 0.045 |
| Arcn1 | Coatomer subunit delta | 16 | 8 | 1 | NaN | NaN | NaN | NaN | 3 | -3.38 | 0.048 |
| Mapk8ip3 | C-Jun-amino-terminal kinase-interacting protein 3 | 11 | 14 | NA | NaN | 2 | NaN | NaN | 5 | -3.23 | 0.048 |
| Rab31 | Ras-related protein Rab-31 | 12 | 6 | 1 | NaN | NaN | NaN | NaN | 1 | -3.22 | 0.049 |
| Hip1r | Huntingtin-interacting protein 1-related protein | 10 | 8 | NA | NaN | NaN | NaN | 3 | NaN | -3.13 | 0.052 |
| Snx18 | Sorting nexin | 7 | 11 | NA | NaN | 1 | NaN | NaN | 2 | -3.10 | 0.053 |
| Uso1 | General vesicular transport factor p115 | 8 | 6 | NA | 1 | NaN | NaN | NaN | NaN | -2.91 | 0.053 |
| Snx3 | Sorting nexin-3 | 6 | 7 | NA | NaN | NaN | NaN | NaN | 3 | -2.85 | 0.046 |
| Rab5b | Ras-related protein Rab-5B | 5 | 4 | NA | 1 | 1 | NaN | NaN | NaN | -2.75 | 0.054 |
| Sh3glb1 | Isoform 2 of Endophilin-B1 | 4 | 5 | NaN | NaN | NaN | NaN | NaN | NaN | -2.74 | 0.050 |
| Scfd1 | Sec1 family domain-containing protein 1 | 6 | 5 | NA | NaN | 1 | NaN | NaN | 2 | -2.57 | 0.045 |
| Picalm | Phosphatidylinositol-binding clathrin assembly protein | 5 | 3 | NA | NaN | NaN | NaN | NaN | NaN | -2.36 | 0.046 |
| Srcin1 | SRC kinase signaling inhibitor 1 | 20 | 13 | NA | NaN | NaN | NaN | NaN | NaN | -4.57 | 0.055 |
| Hnrnpd | Heterogeneous nuclear ribonucleoprotein D0 | 16 | 11 | NA | NaN | NaN | NaN | NaN | NaN | -4.39 | 0.053 |
| Ampd3 | AMP deaminase 3 | 9 | 6 | NA | NaN | NaN | NaN | NaN | NaN | -4.01 | 0.057 |
| Psmc4 | 26S proteasome regulatory subunit 6B | 10 | 7 | NA | 3 | 2 | NaN | 2 | NaN | -4.01 | 0.046 |
| Nacad | NAC-alpha domain-containing protein 1 | 10 | 6 | NA | NaN | NaN | NaN | 2 | NaN | -4.00 | 0.052 |
| Dip2b | Disco-interacting protein 2 homolog B | 23 | 13 | NA | 1 | 3 | NaN | 1 | 4 | -3.95 | 0.054 |
| Srp68 | Signal recognition particle subunit SRP68 | 14 | 8 | NA | 2 | NaN | NaN | NaN | 1 | -3.93 | 0.048 |
| Ptbp1 | Polypyrimidine tract-binding protein 1 | 19 | 17 | NA | NaN | NaN | NaN | NaN | 4 | -3.79 | 0.055 |
| Map1a | Microtubule-associated protein 1A | 57 | 39 | 4 | 2 | 2 | NaN | 7 | 13 | -3.77 | 0.044 |
| Qki | Isoform 6 of Protein quaking | 20 | 36 | 7 | NaN | NaN | NaN | NaN | NaN | -3.67 | 0.057 |
| Ckap5 | Cytoskeleton-associated protein 5 | 13 | 13 | NA | 1 | 2 | NaN | 4 | 4 | -3.66 | 0.044 |
| Mapre1 | Microtubule-associated protein RP/EB family member 1 | 12 | 10 | 2 | 1 | 1 | NaN | NaN | NaN | -3.65 | 0.052 |
| Rps5 | 40S ribosomal protein S5 (Fragment) | 12 | 7 | 3 | NaN | NaN | NaN | NaN | NaN | -3.60 | 0.045 |
| Hyou1 | Hypoxia up-regulated protein 1 | 30 | 14 | 12 | 5 | 4 | NaN | NaN | NaN | -3.56 | 0.050 |
| Prpf19 | Pre-mRNA-processing factor 19 | 11 | 12 | NA | 4 | 2 | NaN | NaN | 2 | -3.55 | 0.059 |
| Arf5 | ADP-ribosylation factor 5 | 12 | 6 | NA | NaN | 2 | NaN | NaN | NaN | -3.55 | 0.045 |
| Atp5pb | ATP synthase F(0) complex subunit B1, mitochondrial | 25 | 14 | 13 | 3 | NaN | NaN | NaN | NaN | -3.43 | 0.047 |
| Top1 | DNA topoisomerase 1 | 10 | 7 | NA | NaN | 1 | NaN | 2 | 1 | -3.41 | 0.053 |
| Acadl | Long-chain-specific acyl-CoA dehydrogenase, mitochondrial | 13 | 7 | NA | NaN | NaN | NaN | NaN | 4 | -3.37 | 0.048 |
| Dbt | Lipoamide acyltransferase component of branched-chain alpha-keto acid dehydrogenase complex, mitochondrial | 22 | 12 | 13 | NaN | 1 | NaN | NaN | NaN | -3.37 | 0.047 |
| Ilf2 | Interleukin enhancer-binding factor 2 | 11 | 8 | 1 | 2 | 2 | NaN | 2 | NaN | -3.35 | 0.052 |
| Psmc5 | 26S proteasome regulatory subunit 8 | 11 | 20 | NA | 1 | 3 | NaN | 3 | 6 | -3.34 | 0.047 |
| Rps27 | 40S ribosomal protein S27 | 9 | 9 | NA | NaN | 1 | NaN | 2 | NaN | -3.33 | 0.054 |
| Arhgap35 | Rho GTPase-activating protein 35 | 10 | 10 | NA | 1 | NaN | NaN | 1 | 5 | -3.30 | 0.046 |
| Ecpas | Proteasome adapter and scaffold protein ECM29 | 8 | 5 | NA | 2 | 1 | NaN | NaN | NaN | -3.25 | 0.057 |
| Qars1 | Glutamine--tRNA ligase | 12 | 8 | NA | NaN | 4 | NaN | NaN | 5 | -3.24 | 0.047 |
| Ccar2 | Cell cycle and apoptosis regulator protein 2 | 10 | 6 | 2 | 1 | 3 | NaN | NaN | NaN | -3.18 | 0.051 |
| Dnmt3a | DNA (cytosine-5)-methyltransferase 3A | 7 | 8 | 1 | 1 | 3 | NaN | 2 | 1 | -3.16 | 0.057 |
| Rpl5 | 60S ribosomal protein L5 | 23 | 21 | 3 | 2 | 2 | 1 | 2 | 5 | -3.16 | 0.048 |
| Syncrip | Heterogeneous nuclear ribonucleoprotein Q | 14 | 9 | NA | 2 | 5 | NaN | 6 | 3 | -3.15 | 0.048 |
| Eif3l | Eukaryotic translation initiation factor 3 subunit L | 5 | 7 | NA | NaN | 2 | NaN | NaN | NaN | -3.14 | 0.056 |
| Sik3 | Serine/threonine-protein kinase SIK3 | 6 | 5 | NA | NaN | NaN | NaN | NaN | NaN | -3.12 | 0.056 |
| Ptpn11 | Tyrosine-protein phosphatase non-receptor type 11 | 10 | 8 | NA | 1 | NaN | NaN | NaN | 2 | -3.11 | 0.052 |
| Jmy | Junction-mediating and -regulatory protein | 6 | 3 | NA | 1 | NaN | NaN | 1 | NaN | -3.08 | 0.049 |
| Stt3a | Dolichyl-diphosphooligosaccharide--protein glycosyltransferase subunit STT3A | 9 | 6 | NA | NaN | 1 | NaN | NaN | 1 | -3.07 | 0.045 |
| Eef1b | Elongation factor 1-beta | 5 | 6 | NA | NaN | NaN | NaN | 1 | NaN | -3.07 | 0.058 |
| Usp14 | Ubiquitin carboxyl-terminal hydrolase 14 | 4 | 5 | NA | NaN | 1 | NaN | NaN | NaN | -3.06 | 0.047 |
| Erp44 | Endoplasmic reticulum resident protein 44 | 5 | 4 | NA | NaN | NaN | NaN | NaN | NaN | -3.04 | 0.053 |
| Bzw1 | Basic leucine zipper and W2 domain-containing protein 1 | 7 | 7 | NA | NaN | NaN | NaN | NaN | NaN | -3.04 | 0.055 |
| Ssb | Lupus La protein homolog | 6 | 4 | NA | NaN | NaN | NaN | NaN | NaN | -3.02 | 0.049 |
| Dynlt1 | Dynein light chain Tctex-type 1 | 4 | 6 | NA | 2 | NaN | NaN | 1 | NaN | -2.97 | 0.046 |
| Sh3gl1 | Endophilin-A2 | 4 | 4 | NA | NaN | NaN | NaN | NaN | NaN | -2.97 | 0.051 |
| Vdac3 | Voltage-dependent anion-selective channel protein 3 | 19 | 23 | 4 | 5 | 3 | 1 | 2 | 5 | -2.95 | 0.048 |
| Alg2 | Alpha-1,3/1,6-mannosyltransferase ALG2 | 7 | 6 | NA | NaN | NaN | NaN | NaN | 1 | -2.94 | 0.055 |
| Abhd12 | Lysophosphatidylserine lipase ABHD12 | 9 | 16 | NA | NaN | 2 | NaN | NaN | 7 | -2.91 | 0.046 |
| Agpat4 | 1-acyl-sn-glycerol-3-phosphate acyltransferase delta | 10 | 9 | NA | 3 | 2 | 1 | 3 | 5 | -2.89 | 0.046 |
| Rpl36 | 60S ribosomal protein L36 | 8 | 6 | NA | NaN | NaN | NaN | NaN | NaN | -2.89 | 0.044 |
| Rpl30 | 60S ribosomal protein L30 | 4 | 6 | NA | NaN | 2 | NaN | NaN | NaN | -2.84 | 0.049 |
| Tra2a | Transformer-2 protein homolog alpha | 4 | 4 | NA | NaN | NaN | NaN | NaN | NaN | -2.84 | 0.046 |
| Hectd4 | HECT domain E3 ubiquitin protein ligase 4 | 3 | 4 | NA | NaN | NaN | NaN | NaN | NaN | -2.83 | 0.044 |
| Pds5b | Sister chromatid cohesion protein PDS5 homolog B | 4 | 3 | NA | NaN | NaN | NaN | NaN | NaN | -2.83 | 0.054 |
| Eif3b | Eukaryotic translation initiation factor 3 subunit B | 12 | 6 | NA | 1 | 1 | NaN | NaN | 4 | -2.81 | 0.047 |
| Ppp1r21 | Protein phosphatase 1 regulatory subunit 21 | 7 | 5 | NA | NaN | NaN | NaN | NaN | NaN | -2.80 | 0.047 |
| Slc25a4 | ADP/ATP translocase 1 | 54 | 75 | 10 | 5 | 3 | 16 | 30 | 5 | -2.78 | 0.047 |
| Phldb1 | Pleckstrin homology-like domain family B member 1 | 8 | 4 | NA | 1 | NaN | NaN | NaN | 1 | -2.72 | 0.047 |
| Mtch1 | Mitochondrial carrier homolog 1 | 5 | 3 | NA | NaN | NaN | NaN | NaN | NaN | -2.72 | 0.046 |
| Plxnb3 | Plexin-B3 | 6 | 10 | NA | 2 | 2 | NaN | NaN | 3 | -2.70 | 0.047 |
| Farsa | Phenylalanine--tRNA ligase alpha subunit | 6 | 5 | NA | 1 | 1 | NaN | NaN | 2 | -2.69 | 0.045 |
| Slc25a18 | Mitochondrial glutamate carrier 2 | 8 | 9 | NA | NaN | NaN | NaN | 1 | 2 | -2.66 | 0.047 |
| Tomm34 | Mitochondrial import receptor subunit TOM34 | 7 | 4 | NA | NaN | NaN | NaN | NaN | NaN | -2.65 | 0.049 |
| Pcyt2 | Ethanolamine-phosphate cytidylyltransferase | 5 | 4 | NA | NaN | NaN | NaN | NaN | NaN | -2.64 | 0.048 |
| Rpl27a | 60S ribosomal protein L27a | 8 | 13 | NA | 2 | 2 | 2 | 3 | 4 | -2.63 | 0.047 |
| Gcn1 | eIF-2-alpha kinase activator GCN1 | 23 | 19 | 5 | 2 | 5 | NaN | 2 | 6 | -2.63 | 0.046 |
| Mvk | Mevalonate kinase | 5 | 3 | NA | NaN | NaN | NaN | NaN | NaN | -2.62 | 0.046 |
| Tubb3 | Tubulin beta-3 chain | 4 | 7 | NA | 2 | NaN | NaN | 3 | 3 | -2.61 | 0.049 |
| Cyp20a1 | Cytochrome P450 20A1 | 4 | 3 | NA | NaN | NaN | NaN | 2 | NaN | -2.60 | 0.046 |

| | | | | | | | | | | | | | | |
|---|---|---|---|---|---|---|---|---|---|---|---|---|---|---|
| Rdh11 | Retinol dehydrogenase 11 | 6 | 8 | | 3 | 3 | 2 | | NaN | 3 | 1 | | -2.54 | 0.047 |
| Rpl24 | 60S ribosomal protein L24 | 14 | 8 | | 3 | 2 | 3 | | 1 | 2 | 2 | | -2.54 | 0.047 |
| Sbf1 | Myotubularin-related protein 5 | 8 | 4 | | NA | NaN | NaN | | NaN | NaN | 3 | | -2.53 | 0.049 |
| Sec23ip | SEC23-interacting protein | 5 | 4 | | NA | NaN | NaN | | NaN | NaN | NaN | | -2.52 | 0.048 |
| Ppme1 | Protein phosphatase methylesterase 1 | 4 | 4 | | NA | NaN | 1 | | NaN | NaN | 1 | | -2.52 | 0.048 |
| Sec63 | Translocation protein SEC63 homolog | 4 | 7 | | NA | NaN | NaN | | NaN | NaN | NaN | | -2.51 | 0.046 |
| Hsdl2 | Hydroxysteroid dehydrogenase-like protein 2 | 6 | 5 | | NA | NaN | NaN | | NaN | NaN | 2 | | -2.49 | 0.047 |
| Slc14a1 | Urea transporter 1 | 5 | 4 | | NA | NaN | 1 | | NaN | NaN | NaN | | -2.48 | 0.047 |
| Il1rap | Isoform 3 of Interleukin-1 receptor accessory protein | 4 | 4 | | NA | NaN | NaN | | NaN | NaN | NaN | | -2.42 | 0.045 |
| Arpc1b | Actin-related protein 2/3 complex subunit 1B | 5 | 5 | | 1 | 2 | NaN | | NaN | 2 | 1 | | -2.42 | 0.047 |
| Gltp | Glycolipid transfer protein | 4 | 4 | | NA | NaN | NaN | | 1 | NaN | 1 | | -2.37 | 0.046 |
| Prkcq | Protein kinase C theta type | 5 | 3 | | NA | NaN | NaN | | NaN | NaN | NaN | | -2.32 | 0.049 |
| Arl2 | ADP-ribosylation factor-like protein 2 | 4 | 3 | | NA | NaN | NaN | | NaN | NaN | NaN | | -2.28 | 0.047 |
| Ranbp3 | Ran-binding protein 3 | 2 | 2 | | NA | NaN | NaN | | NaN | NaN | NaN | | -2.25 | 0.047 |
| Eif4a2 | Eukaryotic initiation factor 4A-II | 5 | 4 | | NA | 2 | NaN | | NaN | 1 | 2 | | -2.22 | 0.049 |
| Arpc4 | Actin-related protein 2/3 complex subunit 4 | 10 | 6 | | 2 | 2 | 2 | | NaN | 2 | 2 | | -2.21 | 0.049 |
| Atxn2 | Ataxin-2 | 4 | 3 | | NA | NaN | NaN | | NaN | NaN | NaN | | -2.21 | 0.045 |
| Mapt | Microtubule-associated protein tau | 3 | 3 | | 1 | NaN | NaN | | NaN | 1 | NaN | | -2.20 | 0.047 |
| Dstn | Destrin | 3 | 3 | | NA | NaN | NaN | | NaN | NaN | NaN | | -2.19 | 0.046 |
| Ddx19a | ATP-dependent RNA helicase DDX19A | 6 | 5 | | NA | NaN | NaN | | NaN | NaN | NaN | | -2.09 | 0.046 |
| Ahcy | Adenosylhomocysteinase | 4 | 3 | | NA | NaN | NaN | | NaN | NaN | 1 | | -2.09 | 0.049 |
| Eif3h | Eukaryotic translation initiation factor 3 subunit H | 3 | 4 | | NA | NaN | 1 | | NaN | NaN | 1 | | -2.07 | 0.049 |
| Agpat1 | 1-acyl-sn-glycerol-3-phosphate acyltransferase alpha | 4 | 3 | | NA | NaN | NaN | | NaN | NaN | 1 | | -2.07 | 0.049 |
| Gnpat | Dihydroxyacetone phosphate acyltransferase | 4 | 3 | | NA | NaN | NaN | | NaN | NaN | 1 | | -2.06 | 0.049 |
| Auh | Methylglutaconyl-CoA hydratase, mitochondrial | 3 | 3 | | NA | NaN | NaN | | NaN | 1 | 1 | | -2.02 | 0.049 |

**Supplementary Table 5:** Mass spectrometry peptide counts for control vs. iBot;*Cnp*-Cre oligodendrocytes

| | Gene | $Log_2$ fold change (Ribo-seq of remyelination) | $-Log_2$ fold depletion iBot vs control mass spectrometry |
|---|---|---|---|
| 1 | Mag | 2.72 | 2.98 |
| 2 | Mbp | 2.62 | 4.05 |
| 3 | Tppp | 1.80 | 3.01 |
| 4 | Nacad | 1.75 | 4.00 |
| 5 | Agpat4 | 1.72 | 2.89 |
| 6 | Plxnb3 | 1.52 | 2.70 |
| 7 | Phldb1 | 1.44 | 2.72 |
| 8 | Sbf1 | 1.43 | 2.53 |
| 9 | Prkcq | 1.43 | 2.32 |
| 10 | Ank3 | 1.41 | 3.82 |
| 11 | Srcin1 | 1.37 | 4.57 |
| 12 | Bin1 | 1.36 | 2.90 |
| 13 | Pcyt2 | 1.32 | 2.64 |
| 14 | Gltp | 1.30 | 2.37 |
| 15 | Mapt | 1.21 | 2.20 |
| 16 | Mvk | 1.10 | 2.62 |
| 17 | Il1rap | 0.98 | 2.42 |
| 18 | Nfasc | 0.97 | 4.90 |
| 19 | Arl2 | 0.93 | 2.28 |
| 20 | Rdh11 | 0.87 | 2.54 |
| 21 | Hip1r | 0.77 | 3.13 |
| 22 | Ptpn11 | 0.70 | 3.11 |
| 23 | Dip2b | 0.68 | 3.95 |
| 24 | Ppp1r21 | 0.66 | 2.80 |
| 25 | Rtn4 | 0.66 | 3.18 |
| 26 | Rab31 | 0.66 | 3.22 |
| 27 | Dst | 0.62 | 4.70 |
| 28 | Dnmt3a | 0.59 | 3.16 |
| 29 | Cdk5 | 0.56 | 2.49 |
| 30 | Sh3glb1 | 0.53 | 2.74 |
| 31 | Sept8 | 0.53 | 2.81 |
| 32 | Auh | 0.49 | 2.02 |
| 33 | Bzw1 | 0.47 | 3.04 |
| 34 | Ilf2 | 0.46 | 3.35 |
| 35 | Ptbp1 | -0.66 | 3.79 |
| 36 | Slc14a1 | -0.83 | 2.48 |

**Supplementary Table 6**: Mass spectrometry hits with differential gene expression from Ribo-seq of a mouse model of remyelination (Voskuhl *et al. PNAS* 2019)
