## Supplementary Figures 1-13 for "CNS myelination requires VAMP2/3-mediated membrane expansion in oligodendrocytes"

**Supplementary Fig. 1**

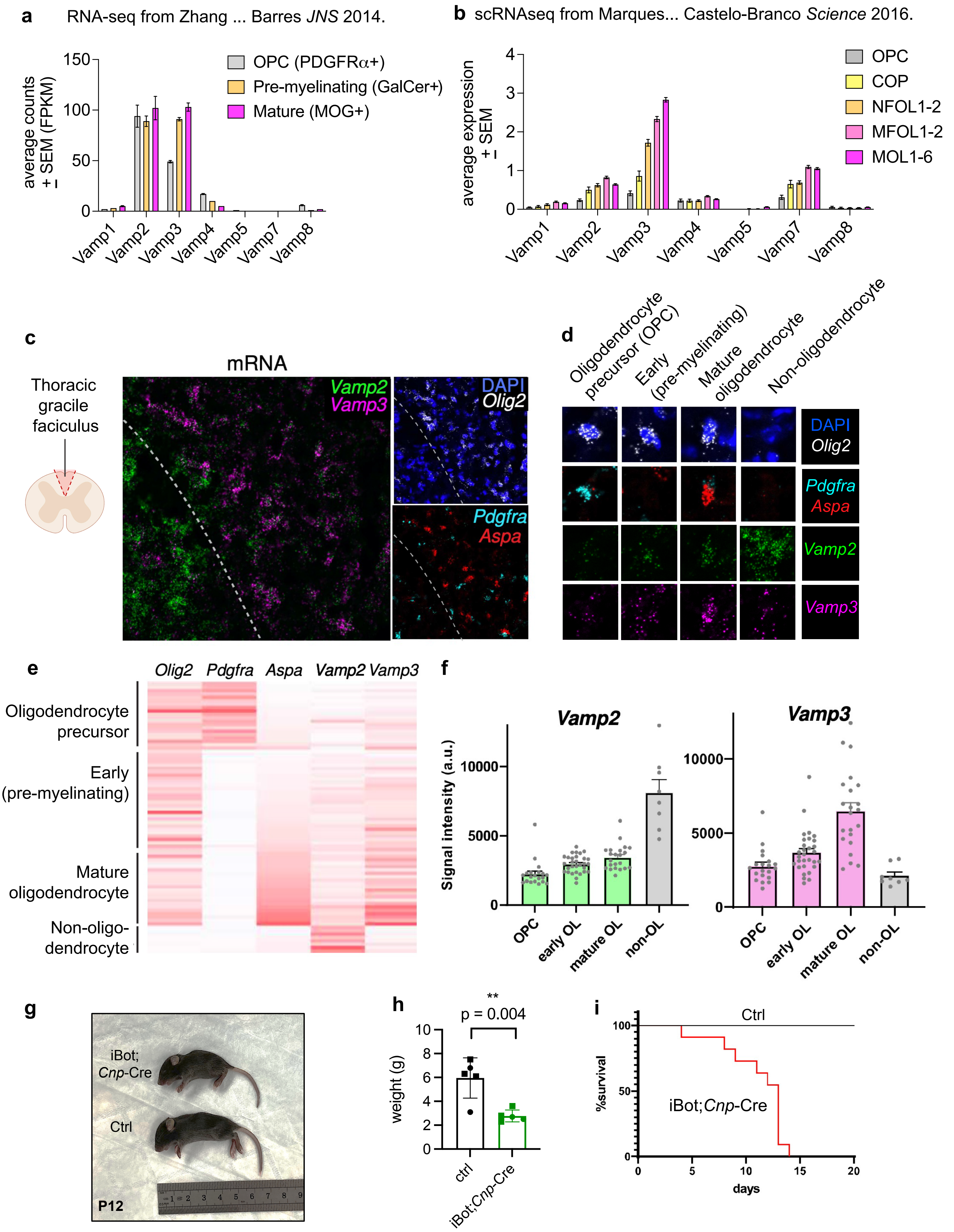

**Supplementary Fig. 1 | Expression of VAMP isoforms in myelinating oligodendrocytes.**

(**a-b**) Expression of v-SNAREs (*Vamp*) in oligodendrocyte-lineage cells from published data sets. (a) Replotted RNA-seq analysis from Zhang *et al. JNS* 2014 with PDGFR$\alpha$+ to identify oligodendrocyte precursors (OPC), GalCer+ to identify pre-myelinating, and MOG+ to mark mature oligodendrocytes. (b) Replotted single cell RNA-seq (scRNA-seq) analysis from Marques *et al*. *Science* 2016, where OPC = oligodendrocyte precursors; COP = differentiation-committed oligodendrocyte precursors; NFOL = newly formed oligodendrocytes; MFOL = myelin-forming oligodendrocytes; MOL = mature oligodendrocytes.

(**c**) Fluorescent *in situ* hybridization (RNAscope) of P10 wild-type mouse spinal cord cross section with probes against the mRNAs *Vamp2*, *Vamp3*, *Olig2, Pdgfra* and *Aspa*. Dotted lines show the boundary between gray matter (left) and white matter (right, dorsal column).

(**d**) Representative images of each cell type, classified by the following markers: OPCs as *Olig2*+ and *Pdgfra*+ cells; early (pre-myelinating) as *Olig2*+, *Pdgfra*- and low *Aspa*; mature oligodendrocytes as *Olig2*+, *Pdgfra*- and high *Aspa* cells; non-oligodendrocytes as *Olig2*- cells.

(**e**) Heat map of fluorescence intensity from each probe in (c), where red is the highest intensity and white is the lowest within a probe, and each line corresponds to a single cell.

(**f**) Bar graph of mRNA fluorescence intensity of *Vamp2* and *Vamp3* in for each cell type. Each datapoint represents an individual cell.

(**g**) iBot;*Cnp*-Cre mouse and control (ctrl) littermate at P12 with ruler showing cm for scale.

(**h**) Body weight (mean $\pm$ SEM) at post-natal day 10 for control littermates (5.79 $\pm$ 0.84 g) and iBot;*Cnp*-Cre mice (2.75 $\pm$ 0.15 g), where squares denote males and circles indicate female with n = 5 for each genotype.

(**i**) Survival curve of iBot;*Cnp*-Cre mice (n = 11) and control littermates (n = 21).

Supplementary Fig. 2

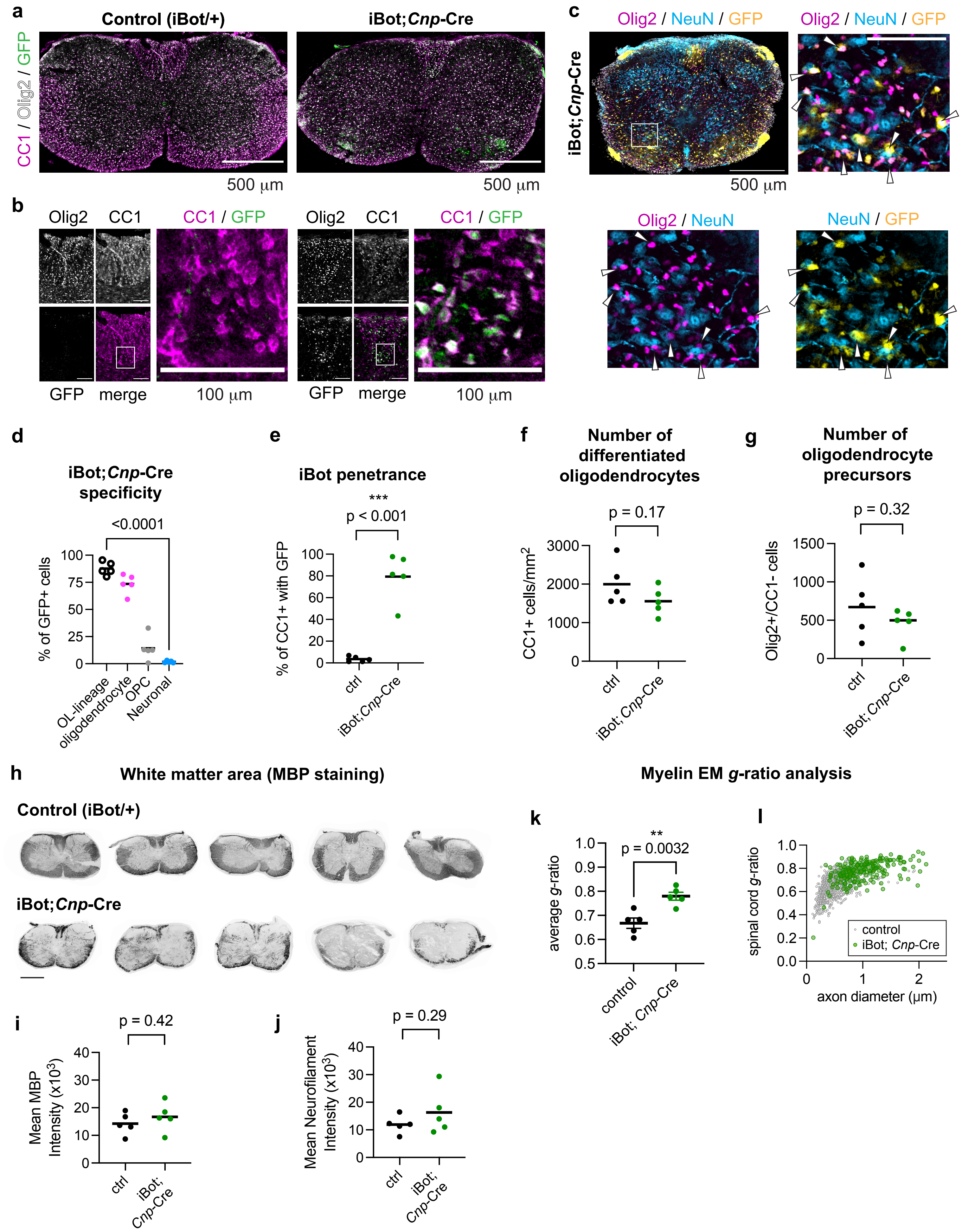

**Supplementary Fig. 2 | Specificity and penetrance of botulinum toxin (iBot) expression in myelinating oligodendrocytes.**

(**a-b**) Immunolabeling of P12 spinal cord cross sections from control (left) and iBot;*Cnp*-Cre (right) mice for CC1 (magenta), Olig2 (white), and GFP to mark iBot expression (green). Scale bar, 500 μm. (b) depicts the dorsal column with enlarged insets to show individual CC1+ cells. Scale bar, 100 μm.

(**c**) Immunolabeling of P12 spinal cord cross sections from iBot;*Cnp*-Cre mice for Olig2 (magenta), NeuN (cyan), and GFP to mark iBot expression (yellow). Scale bar, 500 μm. Inset enlarges a region of the ventral horn, and white arrows point to Olig2+;GFP+ cells. Scale bar, 100 μm.

(**d**) Quantification of iBot;*Cnp*-Cre specificity by counting the % of total GFP+ overlapping with different cell markers (average ± SEM): Olig2 (oligodendrocyte-lineage, 87.8 ± 2.71%), Olig2 and CC1 (oligodendrocyte, 73.5 ± 4.00%), Olig2 without CC1 (OPC, 14.2 ± 5.20%), or NeuN (neuronal, 1.75 ± 0.435%). n = 5 biological replicates with 1067-2085 GFP+ cells analyzed in each.

(**e**) Quantification of iBot penetrance by counting the % of total CC1+ overlapping with GFP (average ± SEM) in control (3.5 ± 1.1%) and iBot;*Cnp*-Cre (79.4 ± 9.7%) littermates for n = 5 biological replicates of each genotype with 1096-2881 CC1+ cells analyzed in each.

(**f**) Quantification of the number of CC1+ cells per $mm^2$ (average ± SEM) for control (1999 ± 249) and iBot;*Cnp*-Cre (1558 ± 160) littermates with n = 5 biological replicates for each genotype.

(**g**) Quantification of the number of CC1-/Olig2+ cells per $mm^2$ (average ± SEM) for control (672 ± 176) and iBot;*Cnp*-Cre (463 ± 87.4) littermates with n = 5 biological replicates for each genotype.

(**h**) Biological replicates of MBP immunolabeling of P12 spinal cord cross sections from control (top row) and iBot;*Cnp*-Cre (bottom row) mice) for the representative image shown in Fig. 1c. Scale bar, 500 μm.

(**i**) Quantification of mean MBP intensity (a.u.) over the entire spinal cord cross section (P12) shown in (h); average $\pm$ SEM for n = 5; control: $(14.2 \pm 1.75) \times 10^3$; iBot;*Cnp*-Cre: $(16.7 \pm 2.31) \times 10^3$).

(**j**) Quantification of mean neurofilament 200 intensity (a.u.) over the entire spinal cord cross section (P12) shown in (h); average $\pm$ SEM for n = 5; control: $(11.9 \pm 1.42) \times 10^3$; iBot;*Cnp*-Cre: $(16.3 \pm 3.58) \times 10^3$).

(**k**) Quantification of *g*-ratio for wrapped/wrapping axons in control vs. iBot;*Cnp*-Cre spinal cords at P12 as shown in Fig. 1f-g; average $\pm$ SEM for n = 5; control: $0.668 \pm 0.0213$; iBot;*Cnp*-Cre: $0.780 \pm 0.0166$).

(**l**) Distribution of *g*-ratio vs. axon diameter for wrapped/wrapping axons in control vs. iBot;*Cnp*-Cre spinal cords at P12 as shown in Fig. 1f-g.

**Supplementary Fig. 3**

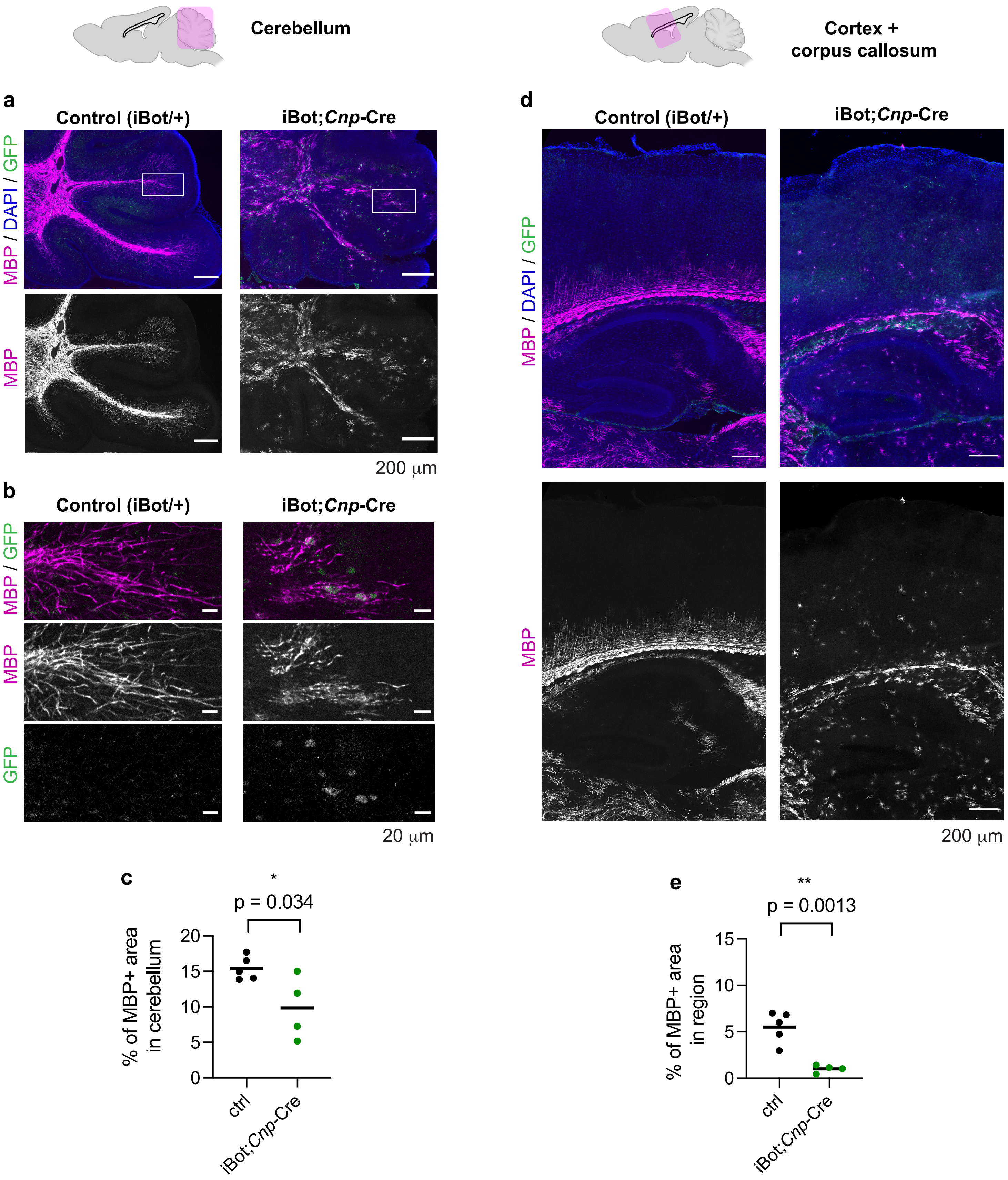

**Supplementary Fig. 3 | VAMP2/3 is required for developmental myelination in the brain.**

(**a**) Immunolabeling of a P12 cerebellum from control (left) and iBot;*Cnp*-Cre (right) mice for MBP (magenta), DAPI (blue), and GFP to mark iBot expression (green). Scale bar, 200 μm.

(**b**) Insets from (a) to show differential MBP morphology of iBot-expressing (GFP+) oligodendrocytes. Scale bar, 20 μm.

(**c**) Quantification of the percent of area immunolabeled by the myelin marker MBP (average ± SEM) across the cerebellum for control (15.4 ± 0.74%, n = 5) and iBot;*Cnp*-Cre (9.8 ± 2.2%, n = 4).

(**d**) Immunolabeling of a sagittal slice of the P12 cortex from control (left) and iBot;*Cnp*-Cre (right) mice for MBP (magenta), DAPI (blue), and GFP to mark iBot expression (green). Scale bar, 200 μm.

(**e**) Quantification of the percent of area immunolabeled by the myelin marker MBP (average ± SEM) across cortical sections for control (5.5 ± 0.75%, n = 5) and iBot;*Cnp*-Cre (1.0 ± 0.2%, n = 4).

### Supplementary Fig. 4

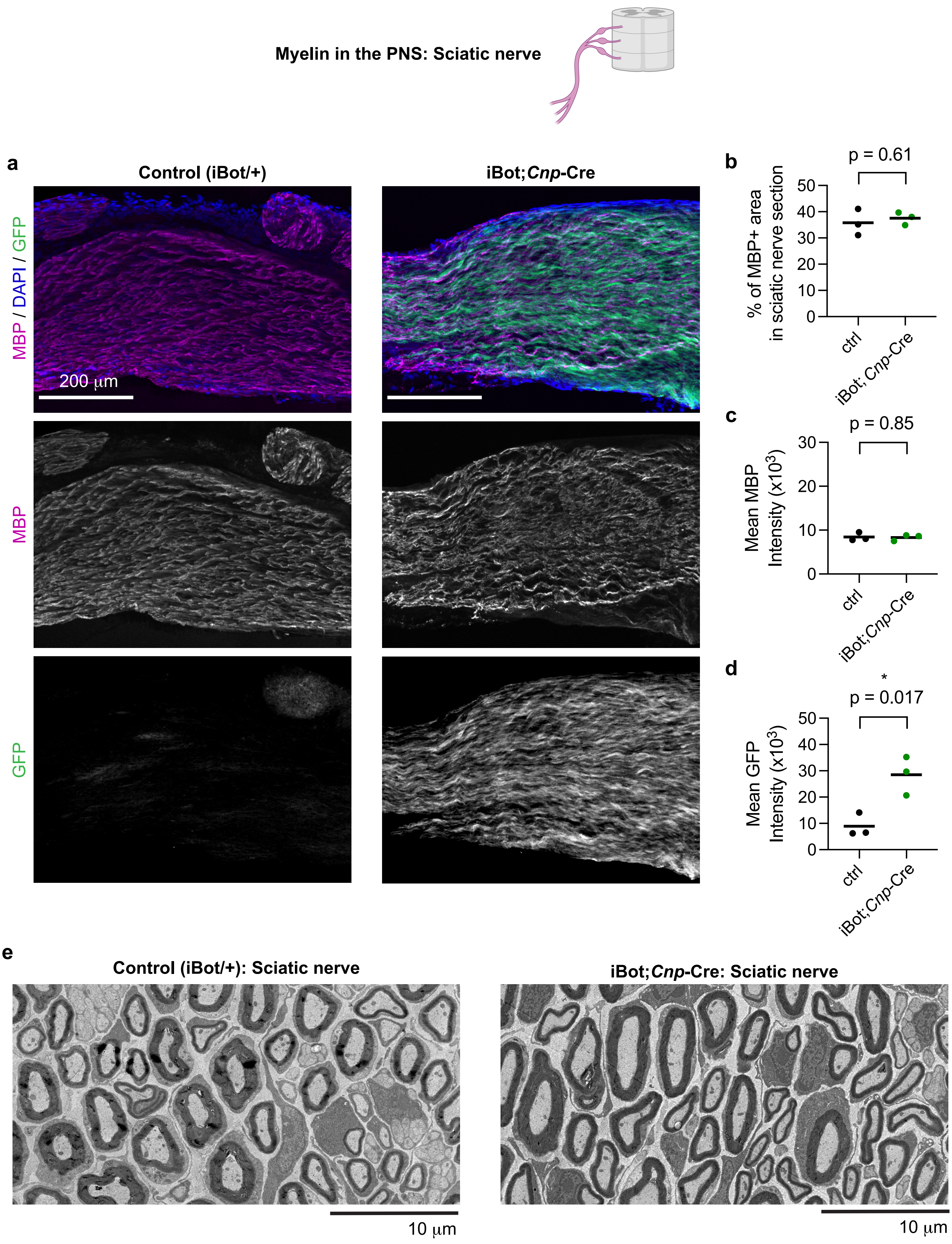

**Supplementary Fig. 4 | Inactivation of VAMP2/3 has no gross effect on PNS myelination.**

(**a**) Immunolabeling of a P12 sciatic nerve longitudinal section from control (left) and iBot;*Cnp*-Cre (right) mice for MBP (magenta), DAPI (blue), and GFP to mark iBot expression (green). Scale bar, 200 μm.

(**b**) Quantification of the percent of area immunolabeled by the myelin marker MBP (average ± SEM) across the sciatic nerve for control (35.8 ± 2.9%, n = 3) and iBot;*Cnp*-Cre (37.6 ± 1.4%, n = 3).

(**c**) Quantification of mean intensity of MBP (average ± SEM, a.u. = arbitrary units) in sciatic nerves for control (8442 ± 550 a.u., n = 3) and iBot;*Cnp*-Cre (8311 ± 407 a.u., n = 3).

(**d**) Quantification of mean intensity of GFP (average ± SEM, a.u. = arbitrary units) in sciatic nerves for control (8942 ± 2599, n = 3) and iBot;*Cnp*-Cre (28534 ± 4244, n = 3).

(**e**) Transmission electron microscopy images from P12 mouse sciatic nerve cross sections (left: control; right: iBot;*Cnp*-Cre). Scale bar, 10 μm.

### Supplementary Fig. 5

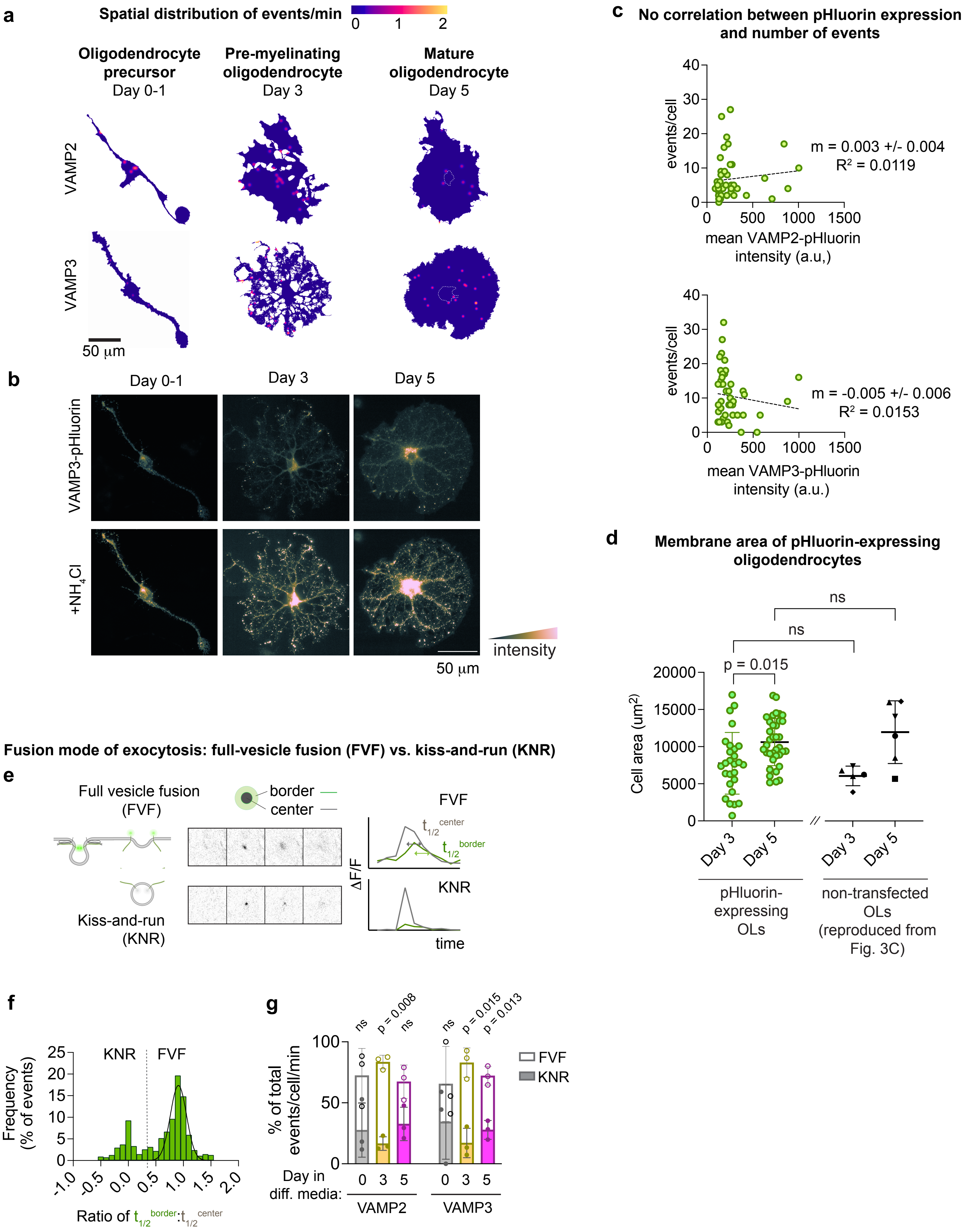

**Supplementary Fig. 5 | Spatial distribution and fusion modes of VAMP2/3-vesicles in primary oligodendrocytes.**

(**a**) Heat map of the density of exocytotic events in primary oligodendrocytes differentiated for 0-1, 3, or 5 days before imaging.

(**b**) Images of primary oligodendrocytes expressing VAMP3-pHluorin (top) followed by a 10-second treatment with 30 mM $NH_4Cl$ (bottom) to de-quench all pHluorin-associated vesicles.

(**c**) Plot of the number of exocytotic events vs. the mean background pHluorin intensity of each cell at t = 0 as a proxy of expression level for VAMP2-pHluorin (top) and VAMP3-pHluorin (bottom). Higher number of exocytotic events did not correlate with higher VAMP-pHluorin expression levels.

(**d**) Comparison of membrane area between pHluorin-expressing (green, pooled data from VAMP2 and VAMP3) and non-transfected primary oligodendrocytes (black) at Day 3 and Day 5 of differentiation in culture. Data shown for non-transfected primary oligodendrocytes were reproduced from the membrane area averages for control oligodendrocytes in Fig. 3c. Mean cell area $\pm$ SEM: at Day 3, pHluorin-expressing 7,761 + 798 $\mu m^2$ (n = 27), non-transfected control 6,099 + 595 $\mu m^2$; at Day 5, pHluorin-expressing 10,607 + 528 $\mu m^2$ (n = 36), non-transfected control 12,022 + 1,731 $\mu m^2$. Statistical significance was determined by a one-way ANOVA with Bonferroni's correction for multiple comparisons.

(**e**) Examples of exocytotic events resulting in different fusion modes of exocytosis. On the top, a full-vesicle fusion (FVF) event exhibits radial spreading of fluorescence from the initial fluorescent punctum. On the bottom, a kiss-and-run (KNR) event appears and disappears without fluorescence spreading. The half-life of the event center ($t_{1/2}^{center}$) measures the duration of initial fluorescent punctum, and the half-life of the bordering event ($t_{1/2}^{border}$) measures the decay of the surrounding membrane fluorescence. In FVF events, the $t_{1/2}^{border}$ (green trace) should be similar to the $t_{1/2}^{center}$ (gray trace), yielding a ratio of half-lives of ~ 1.

(**f**) Frequency distribution of the ratio of half-lives from the bordering event to the center event ($t_{1/2}^{center}$: $t_{1/2}^{border}$) as a measure of radial fluorescence spreading. Ratios of all pHluorin events (n = 396 for VAMP2 and 539 for VAMP3) distributed into two clusters. FVF was assigned to the cluster ~ 1 (mean $\pm$ SD of Gaussian fit for VAMP2: 0.89 $\pm$ 0.14; VAMP3: 0.92 $\pm$ 0.15). Ratios below three standard deviations of the FVF mean (marked by the dotted line) were classified as KNR.

(**g**) Stacked bar graph for composition of events classified as FVF (top open bars and circles) or KNR (bottom filled bars and circles) at Day 0, 3, and 5 of differentiation in culture corresponding to OPC, pre-myelinating, and mature stages, respectively. Statistical significance was determined by a two-way ANOVA between KNR and FVF at each day of differentiation with Bonferroni's correction for multiple comparisons. For statistically significant differences, the mean $\pm$ SEM% of FVF events was 83.4 $\pm$ 4.5% for VAMP2 at Day 3, 82.9 $\pm$ 9.8% for VAMP3 at Day 3, and 72.1 $\pm$ 6.3% for VAMP3 at Day 5 from n = 3 biological replicates.

### Supplementary Fig. 6

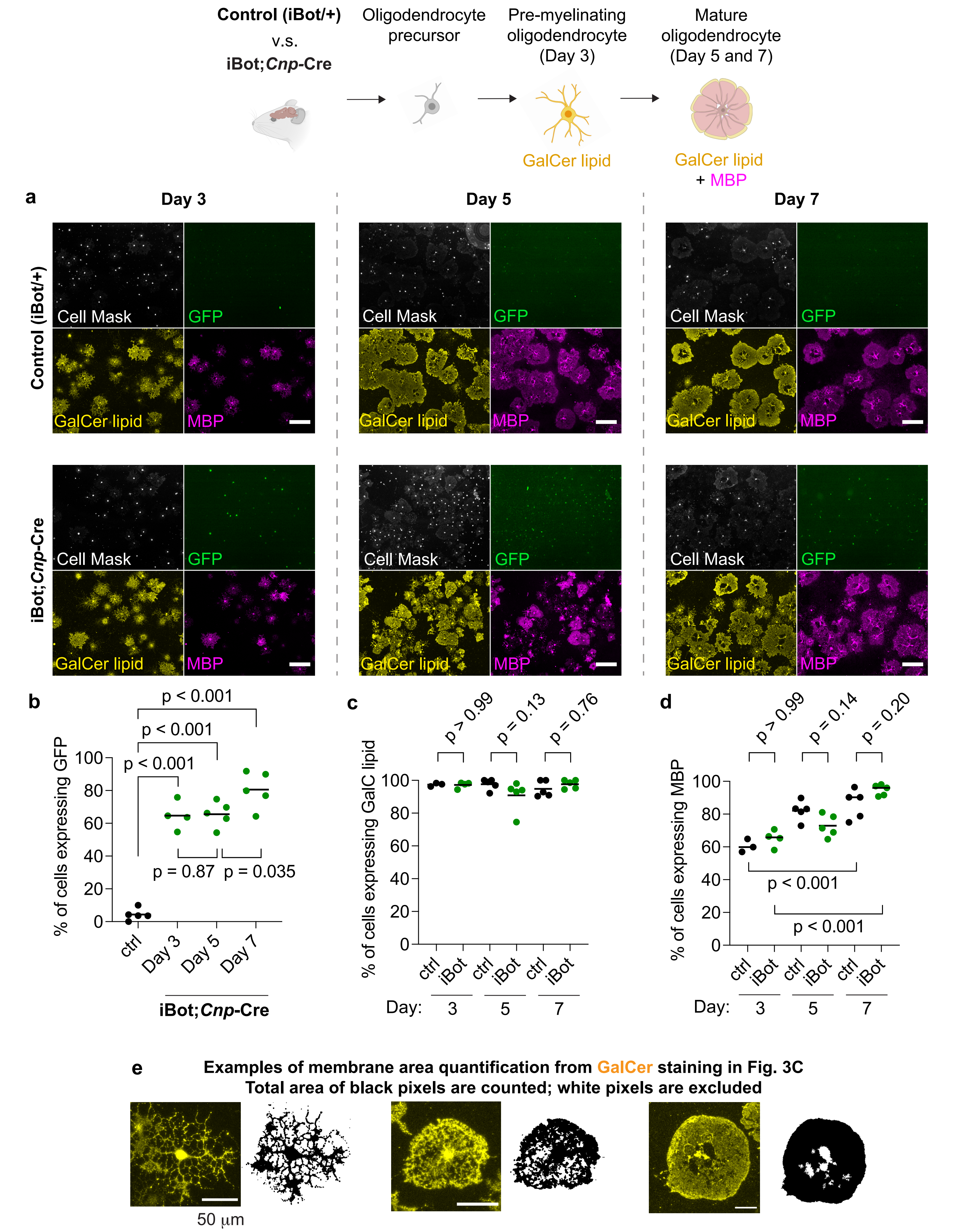

**Supplementary Fig. 6 | VAMP2/3 inactivation in pre-myelinating oligodendrocytes does not affect oligodendrocyte differentiation in culture.**

(**a**) Oligodendrocyte precursors were purified from the brains of control or iBot;*Cnp*-Cre mice and differentiated in culture for 3, 5, or 7 days. Cells were fixed and stained for Cell Mask Blue (a marker of morphology), GalCer lipid (a marker of pre-myelinating oligodendrocytes), and MBP (a marker of mature oligodendrocytes). Scale bar, 200 μm.

(**b**) Quantification of iBot expression as % of cells expressing GFP (mean ± SEM) for n = 5 biological replicates in each condition. Control: 4.3 ± 1.6%. iBot;*Cnp*-Cre Day 3: 64.7 ± 4.3%, Day 5: 65.6 ± 3.4%, Day 7: 80.6 ± 4.9%. Statistical significance was determined by a one-way ANOVA with Bonferroni's correction for multiple comparisons.

(**c**) Quantification of pre-myelinating oligodendrocytes as % of cells expressing GalCer lipid (mean ± SEM) for n = 5 biological replicates in each condition. Control Day 3: 97.4 ± 0.6%, Day 5: 97.6 ± 1.5%, and Day 7: 94.8 ± 2.2%. iBot;*Cnp*-Cre Day 3: 97.2 ± 0.9%, Day 5: 90.9 ± 4.2%, Day 7: 97.7 ± 1.2%. Statistical significance was determined by a one-way ANOVA with Bonferroni's correction for multiple comparisons.

(**d**) Quantification of mature oligodendrocytes as % of cells expressing MBP (mean ± SEM) for n = 5 biological replicates in each condition. Control Day 3: 60.6 ± 2.3%, Day 5: 82.1 ± 2.7%, and Day 7: 86.2 ± 4.0%. iBot;*Cnp*-Cre Day 3: 65.1 ± 2.6%, Day 5: 72.9 ± 3.0%, Day 7: 94.7 ± 1.5%. Statistical significance was determined by a one-way ANOVA with Bonferroni's correction for multiple comparisons.

(**e**) Examples of membrane area quantification from GalCer staining in Fig. 3C to show how unstained regions arising from the reticulated membrane network are excluded from area quantification.

### Supplementary Fig. 7

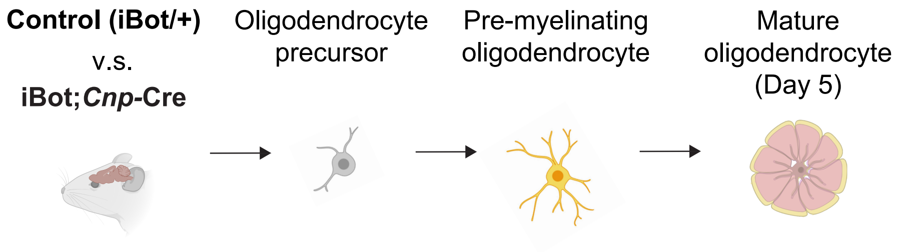


**a**

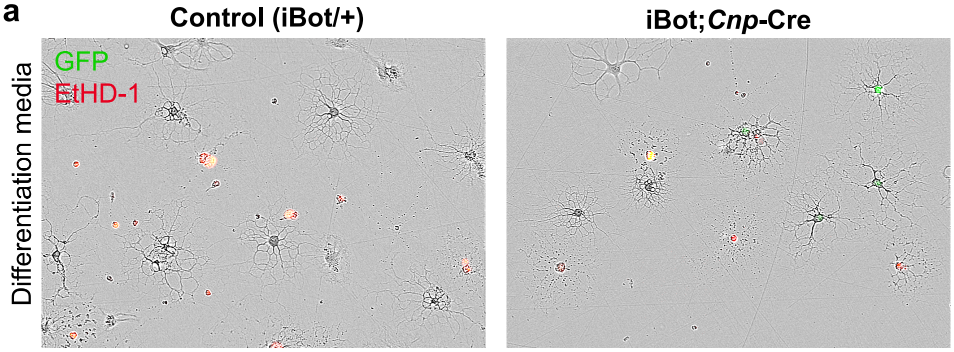


**b**

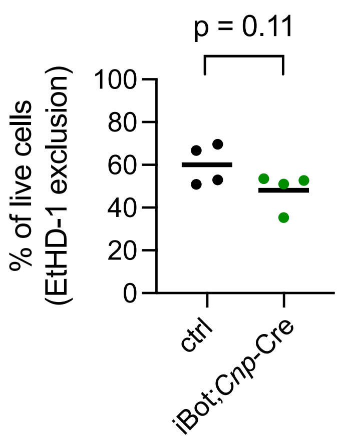


**c**

| Genotype | Rep | Total cells | EtHD-1+ | %EtHD-1+ | %live |
|---|---|---|---|---|---|
| ctrl | 1 | 194 | 59 | 30% | 70% |
| | 2 | 171 | 57 | 33% | 67% |
| | 3 | 204 | 96 | 47% | 53% |
| | 4 | 165 | 81 | 49% | 51% |
| iBot | 1 | 153 | 71 | 46% | 54% |
| | 2 | 186 | 91 | 49% | 51% |
| | 3 | 245 | 116 | 47% | 53% |
| | 4 | 195 | 126 | 65% | 35% |

**d**

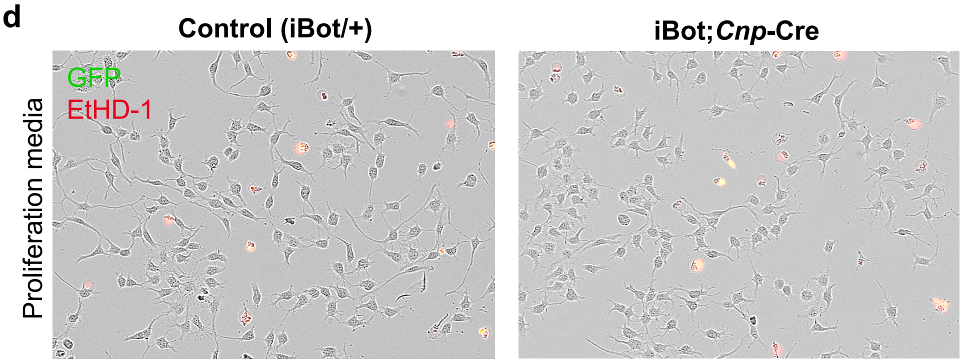


**e**

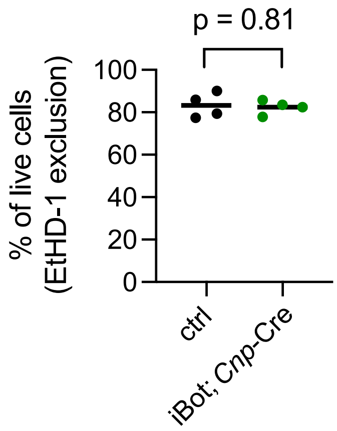


**f**

| Genotype | Rep | Total cells | EtHD-1+ | %EtHD-1+ | %live |
|---|---|---|---|---|---|
| ctrl | 1 | 1597 | 159 | 10% | 90% |
| ctrl | 2 | 1002 | 226 | 23% | 77% |
| ctrl | 3 | 1637 | 230 | 14% | 86% |
| ctrl | 4 | 813 | 168 | 21% | 79% |
| iBot | 1 | 1177 | 167 | 14% | 86% |
| iBot | 2 | 940 | 155 | 16% | 84% |
| iBot | 3 | 1450 | 255 | 18% | 82% |
| iBot | 4 | 675 | 150 | 22% | 78% |

**Supplementary Fig. 7 | VAMP2/3 inactivation does not affect oligodendrocyte cell survival.**

(**a**) Dead cell staining of primary oligodendrocytes at Day 5 in differentiation media with ethidium homodimer (EtHD-1, red).

(**b**) Quantification of live oligodendrocytes as % of cells that excluded ethidium homodimer (mean $\pm$ SEM) with n = 4 biological replicates for control (60.0 $\pm$ 4.7%) and iBot;*Cnp*-Cre (48.2 $\pm$ 4.3%).

(**c**) Number of oligodendrocytes quantified in each biological replicate.

(**d**) Dead cell staining of oligodendrocyte precursors in proliferation media with ethidium homodimer (EtHD-1, red).

(**e**) Quantification of live precursors as % of cells that excluded ethidium homodimer (mean $\pm$ SEM) with n = 4 biological replicates for control (83.2 $\pm$ 2.9%) and iBot;*Cnp*-Cre (82.4 $\pm$ 1.7%).

(**f**) Number of oligodendrocyte precursors quantified in each biological replicate.

### Supplementary Fig. 8

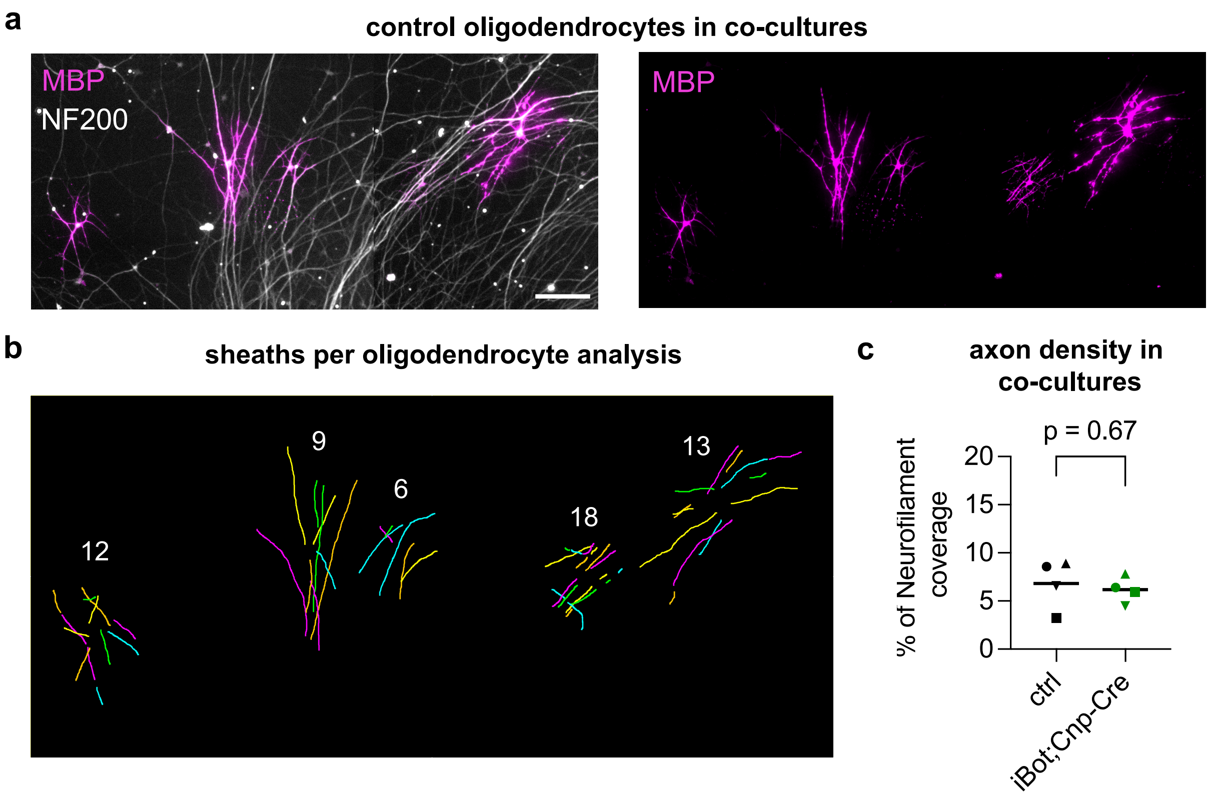

**Supplementary Fig. 8 | Example of sheath number per cell analysis**

(**a**) Primary oligodendrocytes purified from control mouse brain cultured on CNS-derived axons (retinal ganglion cell re-aggregates) for 7 days and stained for MBP (magenta) and NF200 (white). Scale bar 100 $\mu$m.

(**b**) Examples of sheath analysis for individual oligodendrocytes in co-cultures, where each color depicts a distinct sheath. The white number corresponds to the number of sheaths counted per cell.

(**c**) Quantification of axon density by neurofilament-200 staining in co-cultures shown in Fig. 4b-d.

### Supplementary Fig. 9

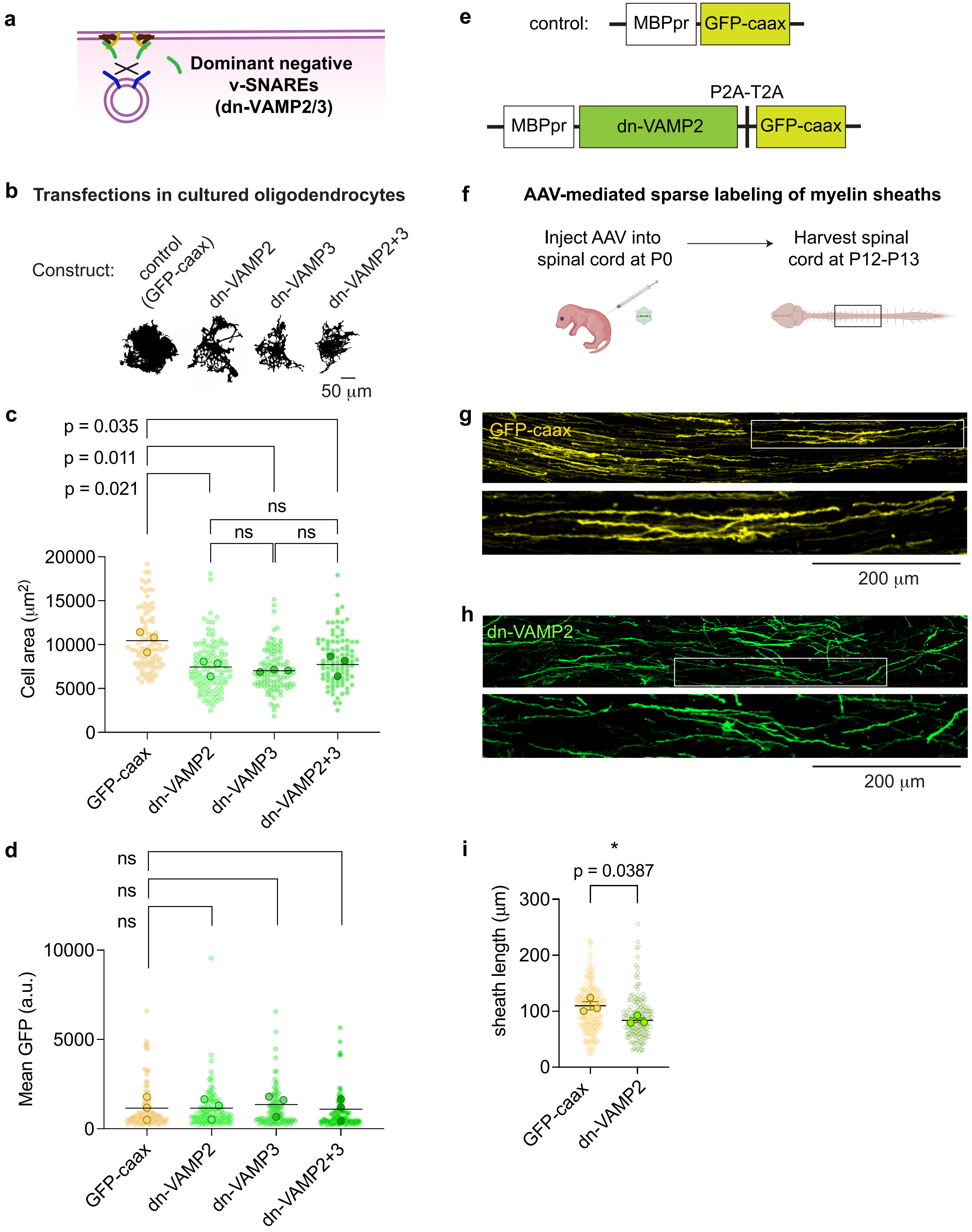

**Supplementary Fig. 9 | Expression of dominant negative VAMP (dn-VAMP2/3) as an orthogonal method of inhibiting VAMP2/3-mediated exocytosis**

(**a**) Schematic of dominant negative v-SNAREs (green, dn-VAMP2/3) inhibiting membrane fusion by sequestering binding sites on the target membrane from endogenous v-SNAREs (blue).

(**b**) GalCer lipid-stained masks of primary rat oligodendrocytes transfected with MBP promoter-driven dn-VAMP2, dn-VAMP3, or both dn-VAMP2 and 3 (dn-VAMP2+3), and differentiated for 5 days in culture to show cell morphologies. Scale bar, 50 μm.

(**c**) Quantification of membrane surface area marked by GalCer lipid, where each light-shaded point corresponds to a single cell, each dark-shaded point represents the mean area of cells from one biological replicate, and the line represents the mean area from n = 3 biological replicates for each condition. Mean cell area ± SEM after Day 5 of differentiation: GFP-caax (control) 10,457 ± 690 $\mu m^2$; dn-VAMP2 7,454 ± 531 $\mu m^2$; dn-VAMP3 7,040 ± 78.5 $\mu m^2$; dn-VAMP2+3 7,745 ± 687 $\mu m^2$. Statistical significance was determined by a one-way ANOVA with Bonferroni's correction for multiple comparisons.

(**d**) Quantification of mean GFP to compare expression levels of dn-VAMP constructs,where each light-shaded point corresponds to a single cell, each dark-shaded point represents the mean area of cells from one biological replicate, and the line represents the mean area from n = 3 biological replicates for each condition. Mean GFP intensity ± SEM after Day 5 of differentiation (a.u.): GFP-caax (control) 1,153 ± 371; dn-VAMP2 1,154 ± 338; dn-VAMP3 1,357 ± 347; dn-VAMP2+3 1,093 ± 356. Statistical significance was determined by a one-way ANOVA with Bonferroni's correction for multiple comparisons.

(**e**) Schematic of constructs used for AAV-mediated sparse labeling of oligodendrocytes, where the control corresponds to MBP promoter (MBPpr)-driven GFP-caax (top), and the experimental condition uses MBPpr-driven dn-VAMP2 followed by a tandem self-cleaving peptide (P2A-T2A) and GFP-caax.

(**f**) Schematic of neonatal mouse pup spinal cord injections for AAV-mediated sparse labeling of oligodendrocytes, where the lumbar spinal cord is injected at P0 and the thoracic spinal cord region is harvested at P12-P13 for sheath analysis.

(**g**) Representative image of oligodendrocytes expressing GFP-caax in the P12 mouse spinal cord. Inset depicts a single, discrete oligodendrocyte and its corresponding sheaths. Scale bar for inset, 200 μm.

(**h**) Representative image of oligodendrocytes expressing dn-VAMP2-P2A-T2A-GFP-caax in the P12 mouse spinal cord. Inset depicts a single, discrete oligodendrocyte and its corresponding sheaths. Scale bar for inset, 200 μm.

(**i**) Quantification of sheath lengths labeled by GFP-caax for control vs. dn-VAMP2, where each light-shaded point corresponds to a single cell, each dark-shaded point represents the mean area of cells from one biological replicate, and the line represents the mean area from n = 3 biological replicates for each condition. Mean sheath length ± SEM: GFP-caax (control) 110 ± 7.31 μm; dn-VAMP2 84.0 ± 4.45 μm.

**Supplementary Fig. 10**

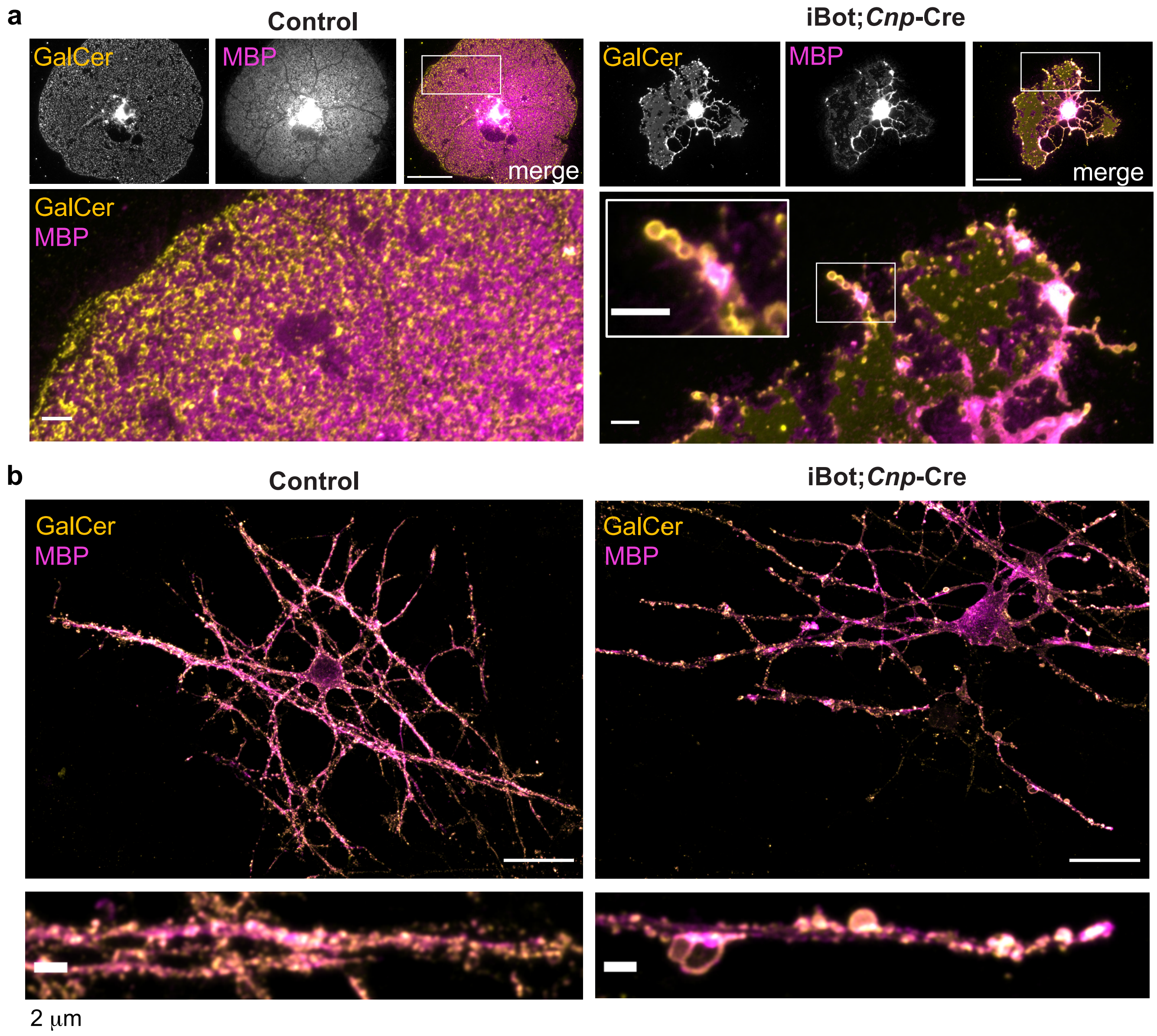

**Supplementary Fig. 10 | VAMP2/3 inactivation induces vesicle accumulation in oligodendrocyte processes and sheaths in culture.**

(**a**) Confocal images of mature oligodendrocytes from control (left) and iBot;*Cnp*-Cre (right) mice stained for GalCer lipid and MBP. Scale bars: top row, 50 μm; bottom row and inset, 5 μm.

(**b**) Confocal images of oligodendrocyte-neuron co-cultures from control (left) and iBot;*Cnp*-Cre (right) oligodendrocytes stained for GalCer lipid and MBP. Scale bars: top row, 20 μm; bottom row and inset, 2 μm.

### Supplementary Fig. 11

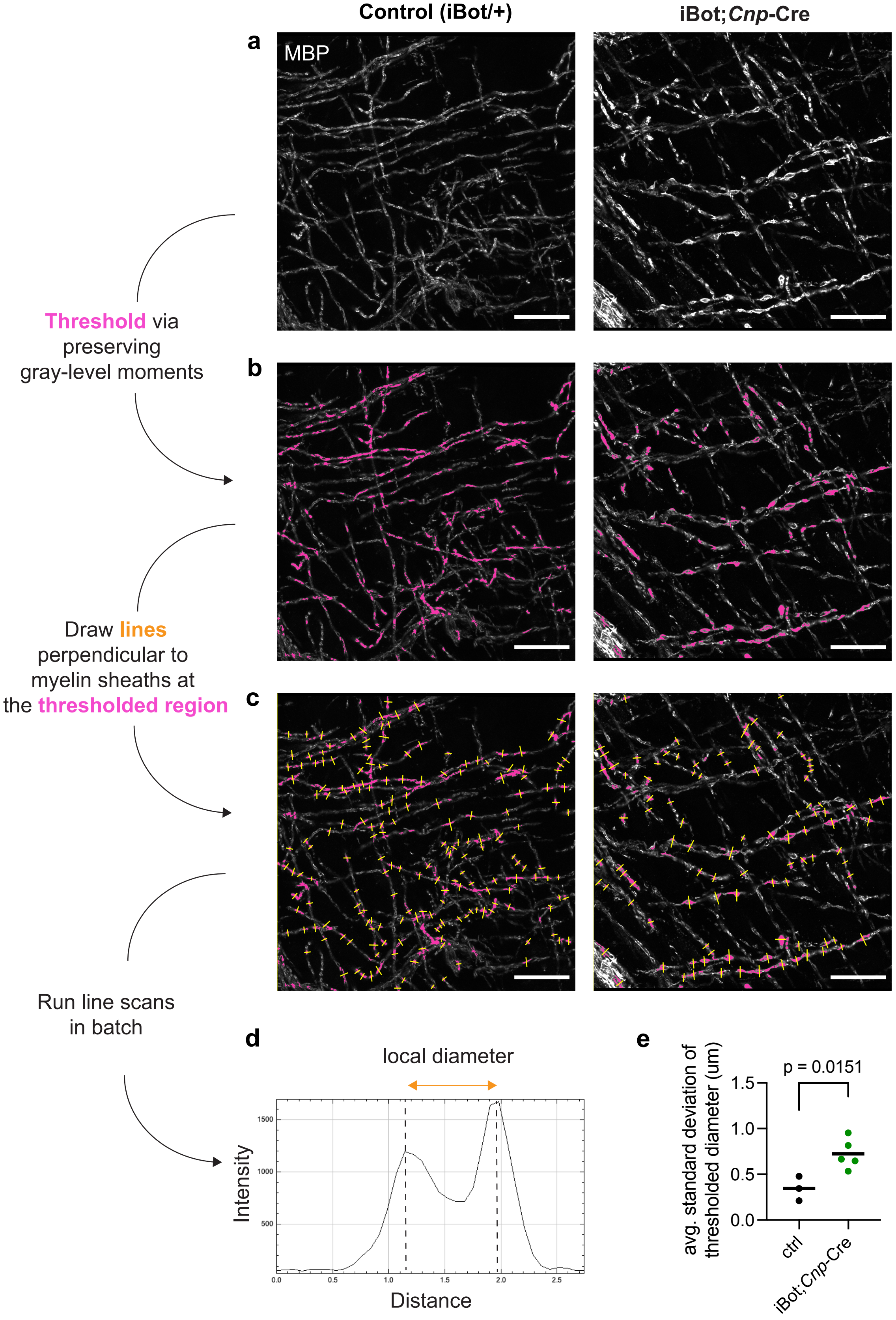

**Supplementary Fig. 11 | Methodology for quantifying local diameter of myelin sheaths in Fig. 5c.**

(**a**) Airyscan confocal images of P12 mouse cingulate cortex from control (left) and iBot;*Cnp*-Cre (right) littermates immunostained for MBP. Scale bar, 20 μm.

(**b**) The pink patches highlight regions thresholded for MBP intensity by preserving gray-level moments (Fiji).

(**c**) Yellow depicts lines that are drawn perpendicular to the longest length of each thresholded pink region. Each yellow line is perpendicular to a myelin sheath.

(**d**) A line scan of each yellow line traces the intensity vs. distance along the line, resulting in two local maxima that correspond to the paired parallel MBP tracks within a sheath. The distance between the two local maxima corresponds to the local diameter of the myelin sheath.

(**e**) The average of the standard deviation (st. dev.) among local diameters measured within a biological replicate. Mean st. dev. $\pm$ SEM: control 0.345 $\pm$ 0.0769 μm from n = 3 biological replicates, iBot;*Cnp*-Cre 0.723 $\pm$ 0.0730 μm from n = 5 biological replicates.

### Supplementary Fig. 12

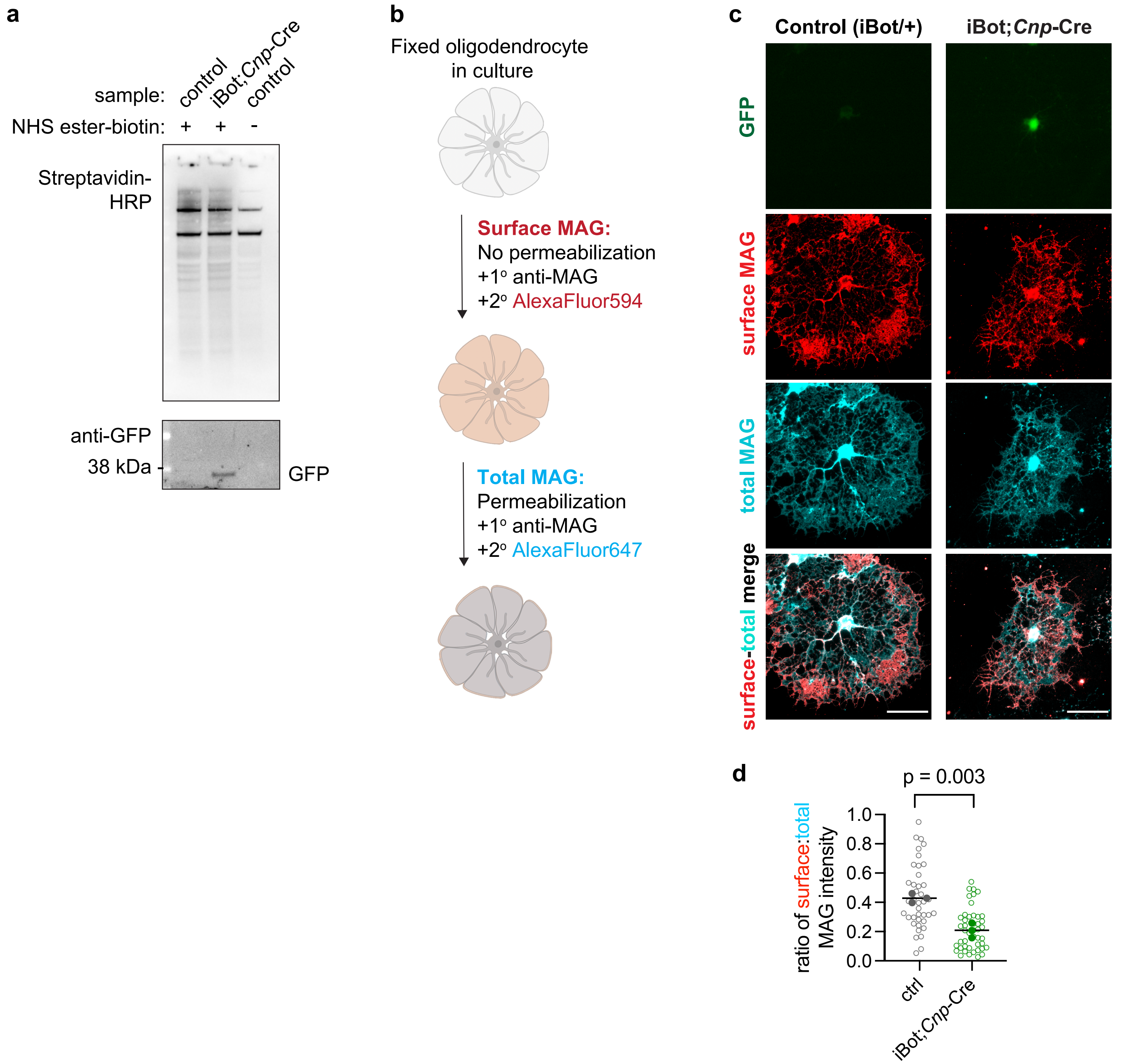

**Supplementary Fig. 12 | Mass spectrometry of surface-biotinylated proteins from oligodendrocytes reveals that the transport of myelin-associated glycoprotein depends on VAMP2/3.**

(**a**) Western blot of mass spectrometry inputs probed for biotinylated substrates (with streptavidin-HRP) and for iBot expression via anti-GFP.

(**b**) Schematic for co-staining cell surface and intracellular myelin associated glycoprotein (MAG), where permeabilization refers to treatment with 0.1% Triton X-100.

(**c**) Representative images of surface and intracellular staining for MAG in control (left) and iBot;*Cnp*-Cre (right) oligodendrocytes. In control cells, surface MAG localizes to the cell periphery, outlining the outer rim of the cell.

(**d**) Quantification of surface MAG intensity relative to total MAG intensity (mean $\pm$ SEM) for control (0.43 $\pm$ 0.02) and iBot;*Cnp*-Cre (0.21 $\pm$ 0.03) from n = 3 biological replicates.

### Supplementary Fig. 13

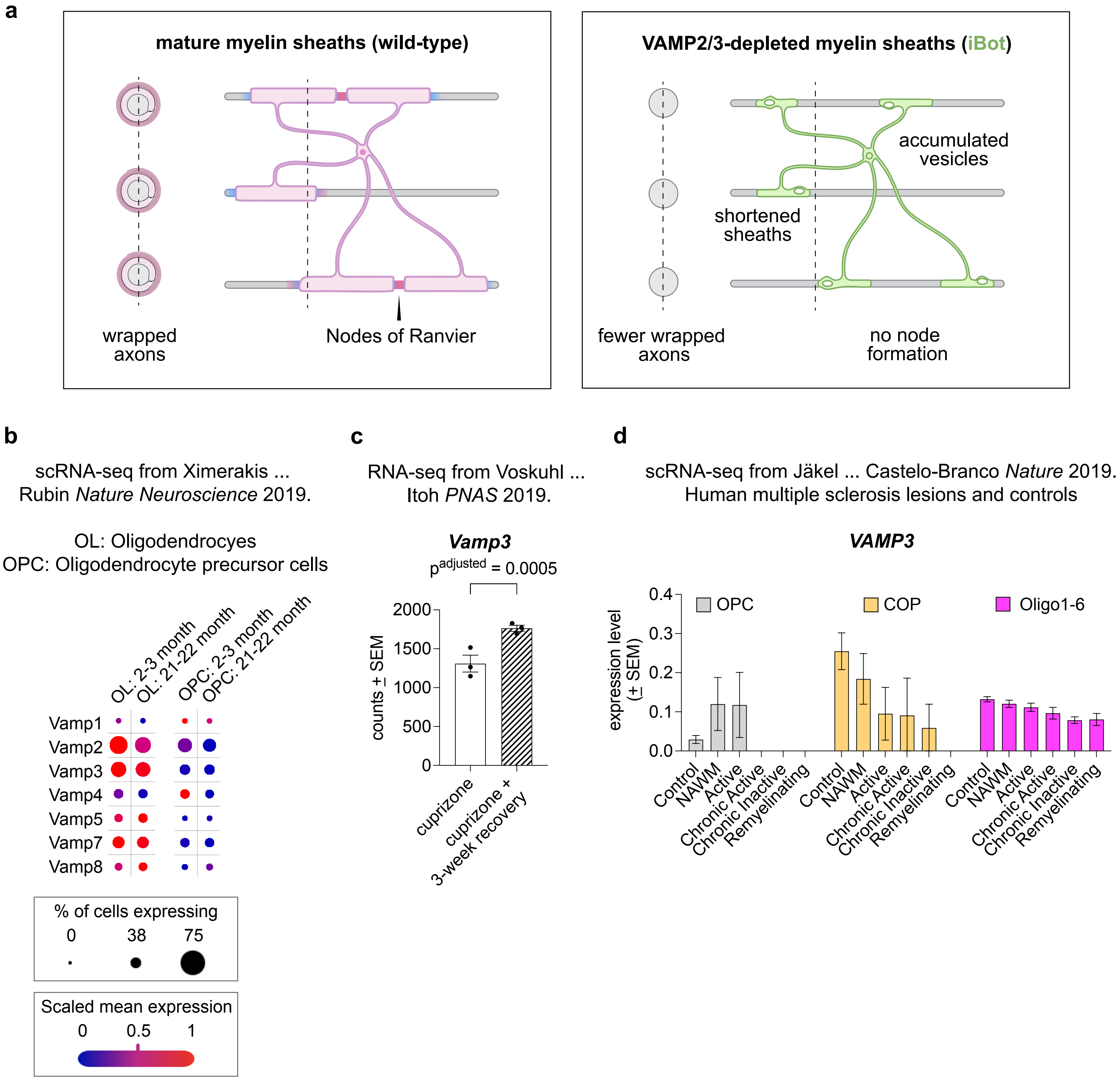

**Supplementary Fig. 13 | VAMP2/3 expression from published datasets of transcriptomic changes in aging and multiple sclerosis.**

(**a**) Depletion of VAMP2/3 by botulinum toxin expression (iBot) causes an accumulation of unfused vesicles in myelin sheaths, preventing sheath elongation and node of Ranvier formation. Shorter sheaths result in reduced myelin coverage of axons, which appears as a higher percentage of unmyelinated axons by TEM. See Methods for a quantitative estimation of relative myelin coverage by control and iBot;*Cnp*-Cre.

(**b**) Dot plot of mean expression of v-SNARE isoforms in oligodendrocytes and OPCs from single-cell RNAseq of adult (2-3 month) and aged (21-22 month) mice published in Ximerakis *et al. Nat. Neuroscien*ce 2019. Plots were generated from the interactive viewer on https://singlecell.broadinstitute.org/single_cell/study/SCP263/aging-mouse-brain.

(**c**) Replotted RNA-seq counts from Voskuhl *et al. PNAS* 2019 for *Vamp3* mRNA associated with oligodendrocyte-derived ribosomes. Samples were isolated from the corpus callosum of mice undergoing demyelination (cuprizone) versus remyelination (cuprizone + 3-week recovery), where $p^{adjusted}$ reports the p-value corrected for false discovery rate.

(**d**) Replotted single cell RNA-seq (scRNA-seq) data from Jakel *et al. Nature* 2019 for *VAMP3* mRNA in white matter from post-mortem tissue of human controls (control) versus patients with progressive multiple sclerosis (MS). The following white matter areas were from MS patients: normal-appearing white matter (NAWM), active, chronic active, chronic inactive, and remyelinating. The different cell types depicted are oligodendrocyte precursors (OPC), differentiation-committed oligodendrocytes (COP), and differentiated oligodendrocytes (Oligo1-6).
